# A Novel Network Approach to Identify Sample-Specific Context-Informed Metabolic Signatures During Developmental Processes

**DOI:** 10.64898/2026.05.20.726642

**Authors:** Emma Lee, Ashwin Koppayi, Almudena Veiga-Lopez, Beatriz Peñalver Bernabé

## Abstract

Metabolism plays an essential role in cellular processes: development, growth, differentiation, and determination of cell identity. Understanding how metabolic processes dynamically change across cell types, stages, and environmental conditions is crucial for studying developmental biology, aging, and disease progression. Genome-wide metabolic models (GEMs) are a powerful network-based tool for studying these processes by integrating omics data to model context-specific metabolism. However, current approaches, such as Flux Balance Analysis (FBA), have limitations in addressing the dynamic nature of metabolism across developmental stages at a sample-specific resolution. To address this, we introduce a novel network-based method for analyzing cell and stage specific metabolic flow using directed and weighted metabolic networks that account for sample-specific transcriptomic data. We apply this method to study ovarian follicle development, providing a deeper understanding of intra-cellular metabolic processes, identifying key metabolites, enzymes, and potential markers for follicular maturation, important for IVF. By incorporating biologically meaningful data, this approach bridges the gap between theoretical metabolic network models (GEMs) and experimental observations, offering a systems-level view of metabolic dynamics in developmental and understudied contexts.

## INTRODUCTION

Metabolism is fundamental during cellular development, supporting critical processes such as growth, proliferation, differentiation, energy production, and establishment of cell identity^1^. Understanding how metabolic processes change over time, between cell types and organs, and across locations within a tissue is essential for better understanding developmental processes, including disease progression or cellular aging.

Systems metabolic approaches have been instrumental for elucidating the complex interplay of inter- and intra-cellular metabolic communications in developmental contexts. Genome-scale metabolic models (GEMs) are network-based tools that allow for the mathematical representation of metabolism in a system (e.g., cell, tissue) and are usually manually curated models that contain known metabolic information of an organism, including reactions and their associated metabolites and the enzymes that catalyze them^2^. By integrating cell-, tissue-, or organ-specific omics data (e.g., transcriptomics, proteomics, metabolomics), GEMs can be tailored to specific biological contexts by constraining the system-wide model to the most likely active metabolic processes^3^. These models have successfully been utilized to predict the metabolic activity of different human tissues, such as liver, brain, heart, and ovarian follicle development in mouse models^4–6^.

However, many computational approaches for studying inter- and intra-cellular metabolism rely heavily on flux balance analysis (FBA), a linear programming-based constrained optimization approach that estimates the optimal metabolic flux distributions under stead-state assumptions to achieve a predefined desirable metabolic output, such as to maximize the production of cellular biomass^7^. While FBA has proven valuable, a significant amount of prior knowledge about the metabolism of the system is required to appropriately define constraints of the system (e.g., metabolic composition of the media around the cells, secretion and consumption rates of crucial metabolites), making the results less insightful in complex and understudied biological contexts, especially *in vivo*^8,9^. Recent efforts to extend network-based analyses beyond FBA have explored graph properties^10–13^. Network methods offer advantages by focusing on structural properties of metabolic networks, such as shortest paths between enzymes (nodes) and cluster (community) identification, providing unique insights into understudied metabolic systems by identifying the hot spots in the metabolism. Importantly, most network characteristics can be determined with little or no prior knowledge of the specific cellular environment. For instance, Beguerisse-Diaz et al. developed a novel approach to identify the most significant metabolic processes and key elements of metabolic communities in *E. coli* and human hepatocyte cells using directed and weighted metabolic networks based on GEMs. The authors determined the directionality of reversible metabolic reactions (whether metabolites are consumed or produced) using the stoichiometric matrix and the FBA results^14^. Directed metabolic networks offer a powerful approach of studying metabolism as they generate more meaningful interpretations of network structures by capturing the directionality of biochemical processes.

However, existing network-based methods face limitations. Many do not incorporate biologically relevant quantitative data from the system being modeled to identify the most significant metabolic processes systematically and in an unbiased manner, nor do they provide a means to compare metabolic activity across individual samples, even though context-specific GEMs can be developed for each individual sample (personalized GEMs). Further, only a few network-based studies explore inter-cellular metabolism in dynamic mammalian developmental processes that include multiple interacting cell types.

To address these challenges, we built on our prior work to derive and integrate context-specific dynamic GEMs to study the progression of ovarian follicle development^6^. Here, we present a novel framework that combines statistical, mathematical, and network analyses methods to explore the dynamic and cell-specific metabolism of complex multi-cellular mammalian developmental processes (**Fig. 1**). We employed human ovarian follicle development as a model system to demonstrate the capabilities of our approach, as human ovarian follicle development is a dynamic and highly coordinated multi-cellular process in which metabolism is crucial^15^. Using publicly available transcriptomic data from two major cell types that comprise the ovarian follicle, the oocyte (female germ-cell) and granulosa cells (somatic cells), at multiple stages during their development (primordial, primary, secondary, and antral stage) and the latest human GEM, Recon3D, we developed the first human ovarian follicle GEMs for each cell type at each stage of development (**Fig. 2**)^16^. We showed that our models provide insights into the potential intra- and inter-cellular metabolic processes that exist in this complex and dynamic process, identify metabolites that may be secreted or consumed during development, and highlight potential biomarkers for ovarian follicle quality and developmental stage, with potential implications for *in vitro* fertilization (IVF).

**Figure 1.**
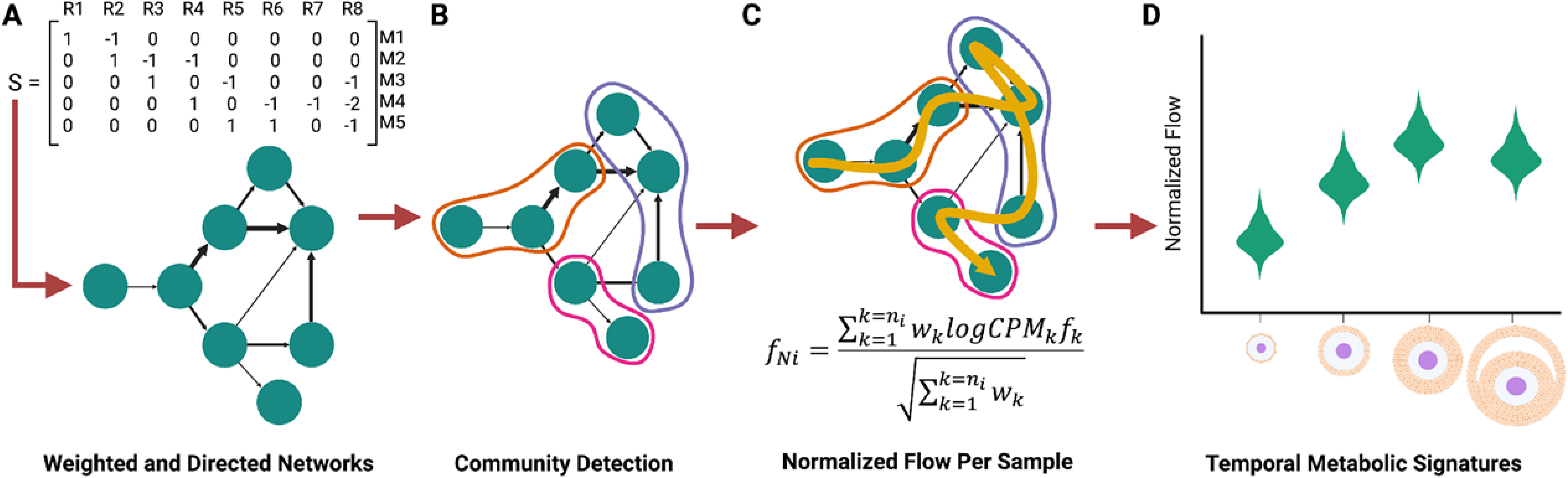
Computational Pipeline to Develop Sample-Specific Metabolic Networks. **A.** The Stoichiometric matrix from a GEM is used to develop a weighted and directed network (which can be be weighted by Flux Balance Analysis results). **B.** Infomap is used for community detection based on flow of information. **C.** The *sample-specific flow* for pathways, enzymes, and metabolites is normalized for each sample (fNi) considering the transcriptional level of each enzyme (logCPM) and the number of metabolites catalyzed by each enzyme (w). **D.** Normalized flows can now be compared across cell types and stages to identify significant temporal metabolic signatures (enzymes, metabolites, pathways).

**Figure 2.**
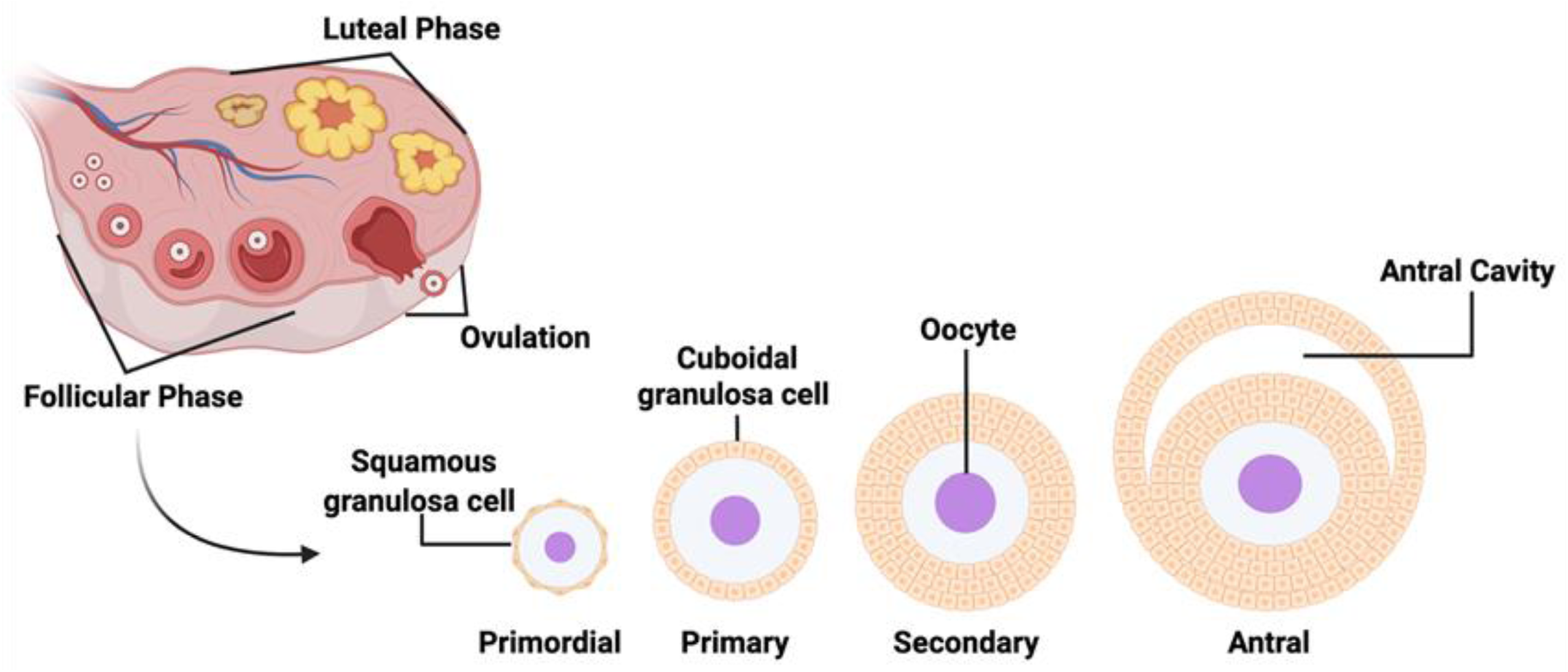
Ovarian Follicle Development. This process begins with the recruitment and activation of primordial follicles from the medulla region of the ovary, which transition into a growing pool of follicles in the cortex. This primordial follicle contains a single layer of squamous granulosa cells surrounding an oocyte and will differentiate into cuboidal granulosa cells to become a primary follicle. Progression to the secondary follicular stage involves granulosa cell proliferation and acquisition of a thecal layer. As antral follicles form, characterized by the development of a fluid-filled cavity, a select few continue to grow to the preovulatory stage, while others undergo atresia.

## METHODS

### Follicle RNA-seq data

We utilized publicly available bulk RNA-seq data of fresh ovarian tissue collected from seven female donors of reproductive age (24 to 32 years old) who underwent an ovariectomy for various reasons (Gene Expression Omnibus (GEO) series GSE107746)^17^. In the study, the authors isolated human follicles from the fresh ovarian tissue and enzymatically digested them to separate the granulosa-oocyte complexes from surrounding stromal and theca cells. Granulosa cells and oocyte cells were separated via enzymatic digestion or aspiration depending on stage, and the zona pellucida of the oocytes was removed^18,19^. Their follicular classification of primordial, primary, secondary, and antral followed Gougeon’s criteria^20^.

The raw paired-end reads from GSE107746 were trimmed and cropped using *Trimmomatic* (v 0.39)^21^. Adapter and Illumina-specific sequences were removed and leading and trailing bases with quality scores below 10 and 25, respectively, were trimmed. Further, starting from the 5’ end, we also trimmed reads whose average quality within a 5-base window fell below 20. Finally, reads shorter than 50 bases after trimming were discarded. We aligned all the processed reads using STAR^22^ to a XX reference genome that was generated by taking the T2T human reference genome and masking the Y chromosome using XYAlign^23–25^. HTSeq-count was used to compute the number of aligned reads^26^.

### Generating genome scale metabolic models

Using the last updated version of Recon3D, with corrections to redox-based reactions identified by Lewis et al.^27^, we generated a follicle GEM with *fastcore.* The inputs for *fastcore* were Recon3D (the global model) and a list of essential reactions determined by the expressed genes that encode for enzymes detected in the samples^16,28^. We defined a gene to be expressed if their weighted ensemble mean expression (EM), a statistically robust measurement against outliers, was greater than log_2_(5) ^29^.

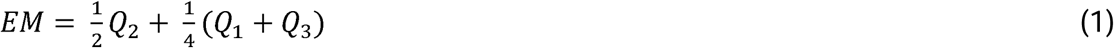

*Q_1_*, *Q_2_*, and *Q_3_*, represent the first, second, and third quartiles of gene expression, respectively. The consistent part of the ovarian follicle GEM was gap filled using the *fastGapFill* function in the Constraint-Based Reconstruction and Analysis Toolbox (COBRA) Toolbox to generate a gap-filled and consistent follicle GEM^30,31^. To generate each cell- and stage-specific models, only expressed genes that encode for enzymes in each cell type at each stage, EM > log_2_(5), were included to determine the core set of reactions. The follicle GEM was used as the global model, instead of Recon3D. Models were constrained using an average plasma composition of females of reproductive age (defined as adult > 18 years old) obtained from the Human Metabolome Database (Table S1) ^32,33^. For each stage, the oocyte and granulosa models were generated. Factoring in the estimated number of granulosa cells per stage based on prior literature (Table S2), granulosa exchange flux values were utilized to update the oocyte models to account for predicted intercellular communication between the cell types^20^. The consistent parts of the models were obtained using the *identifyBlockedReactions* function from the COBRA Toolbox and used for downstream analysis. These constrained models were validated against 481 metabolic functions using the *Test4HumanFct* function in the COBRA Toolbox with the addition of hormone synthesis reactions including the exchange of estradiol, progesterone, and androgen (Table S3).

### Flux balance analysis

FBA is a linear programming approach used to estimate the predicted flow of metabolites through a GEM given a set of constraints and an objective function to optimize its flux^7^. Each cell type at each stage of development was given a different objective function depending on well-known metabolic functions that occur in the cell types at specific stages of development (e.g., estrogen production in granulosa cells), adapted from the mouse ovarian follicle models^6^. For well-studied metabolic processes in the ovarian follicle, such as hormone production, constraints derived from experimental data were placed on corresponding reactions’ lower and upper bounds to ensure biologically meaningful predictions (Table S4). We calculated the predicted flux values for each reaction in each model using the optimizeCbModel from the COBRAToolbox^31^.

### Network construction

Utilizing the methods developed by Beguerisse-Díaz et al., we generated a weighted and directed metabolic network for the follicle model and for each cell- and stage- specific model.^14^ In this format, nodes in the network correspond to reactions, while the directed and weighted edges represent metabolites being transferred from one reaction to another based on the stoichiometric (***S***) matrix, i.e., produced by reaction A and consumed by reaction B. The incorporation of directionality is critical, as it captures the consumption or production of metabolites within biochemical reactions, offering a more precise representation of the directionality of the metabolic connectivity. The first step in generating these networks is to obtain the ***S(m,n)*** matrix for each model with *m* reactions and *n* metabolites. The coefficients within this matrix indicate how much of a metabolite is consumed (-) or produced (+) by a reaction. Since metabolic reactions can be reversible, this method “unfolds” the S matrix into a matrix they define as the ***S_2m_*** matrix, allowing for the explicit representation of forward and reverse reactions. This matrix was constructed as follows:

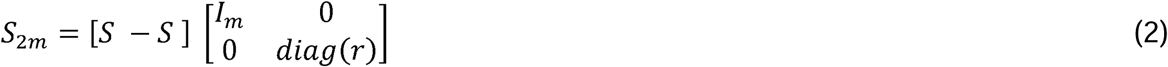

where l*_m_*(*m,m*) is the identity matrix and *diag(r)* is a diagonal *(m,m)* matrix whose diagonal values, *r_j_*, represent reaction reversibility (*r_j_* = 1) or irreversibility (*r_j_* = 1). This transformation doubles the number of reactions in the network, providing a forward and reverse reaction for each reaction. To quantify the predicted metabolite flow between reactions, production 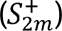 and consumption (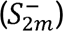) matrices were computed.

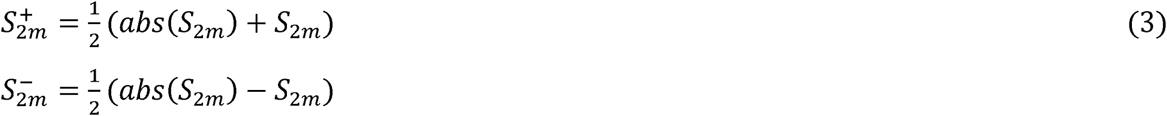

The weight of a given edge between a pair of reactions (nodes), *i* and *j*, can be estimated the probability that a metabolite chosen at random, *k*, is produced by reaction *i* and consumed by a reaction, *j*

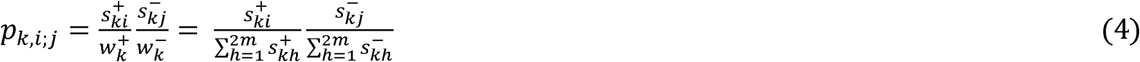

And for all the metabolites in the model, *m*, we can define a normalized edge weight between reactions *i* and *j*, *D_ij_* as:

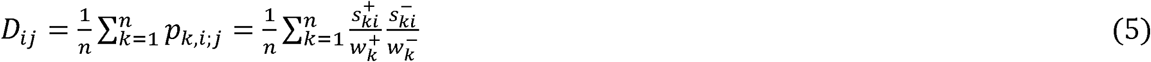

In general, then we can write a normalized edge probability for the entire metabolic network as:

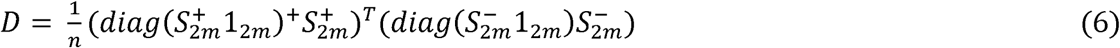

where 1*2m* is a vectors of 1 and + is the Moore Penrose pseudoinverse.

### Community Detection

The physics-derived InfoMap algorithm was utilized for community detection and to model the flow of information for each cell and stage specific metabolic network^34^.

InfoMap utilizes the Map Equation:

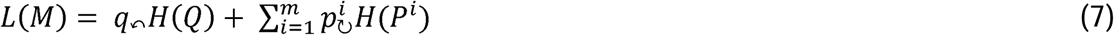

Here, L(M) represents the steps of the random walker for the network (or partition) M. 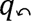 represents the probability of exiting modules, while *H(Q)* represents the entropy of movements between modules.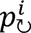 is the probability of using a codebook, i, within a module and *H*(*P^i^*) is the entropy of movements within module i. Together, these terms represent the cost of encoding movements between modules and within them. Minimizing L(M) prioritizes partitions where flow remains within modules. Therefore, InfoMap can identify communities where the random walker stays for longer periods of time before moving to another portion of the network. For each cell and stage specific model, a weighted and directed edge list in plain text format was provided to Infomap. In this list, nodes correspond to reactions from the GEMs and edge weights reflect the weighted importance of metabolic connection as determined by the D matrix. Edge directionality was conserved using the -d option and both flat cluster assignment (--clu) and full hierarchical modular structure (--ftree) were generated. The resulting community partitions and flow values were used in downstream analyses. Communities were defined as the highest community partition. Cytoscape was used to visualize this network structure. The weights of edges between nodes were calculated as the normalized flux of the results between the metabolites that are produced by a reaction and those that are consumed by a reaction across communities. From here, all edges with a weight of 0.001 and above were included in the visualization. Any nodes that had no connections upon this constriction were removed from the visualization. The size of the node corresponds to the number of reactions in the community, and the pie chart within corresponds to the proportion of each subsystem within this community. For visualization purposes the transparency of the edges was adjusted by reducing the transparency on nodes with a weight of less than 0.01.

### Normalized flow analysis

We calculated the flow of information through enzymes, metabolites, metabolic pathways, and communities in the following manner: i) if a reaction could be catalyzed by multiple enzymes, the flow of information of the reaction, calculated using Infomap, was equally split among all associated enzymes—we assumed that the total metabolic flux is the contribution of each associated enzymes independently of their total activity (as it is unknown); ii) if a reaction leads to multiple metabolic products, the flow of information obtained from Infomap was equally split among the reaction products and the flow of information for a given metabolite was the sum of all the reactions that it partakes; iii) the flow of information through a predefined Recon3D pathway was estimated as the summation of the flow of information of each node (reaction) that is part of said metabolic pathway; iv) the flow of information through a community identified by Infomap was the summation of flow of information of all nodes (reactions) within that community. Pathway annotations were assigned based on the Recon3D subsystem classifications. To reduce biases in downstream analysis from highly connected metabolites, the top 2% most common metabolites (e.g., water, ATP) were excluded from the analysis. Next, for each sample, we computed the sample-specific flow values for pathways, enzymes, metabolites, and communities by accounting for the expression level of genes that encode for enzymes. The normalized flow,*f_N_*, for each feature of interest in each sample was determined as:

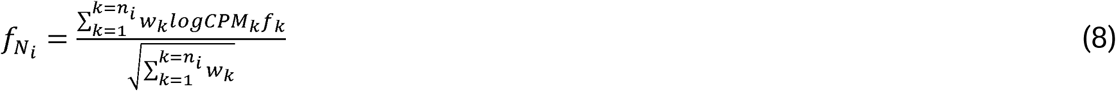

Where *n_i_* is the total number of nodes associate with an enzyme, metabolite, pathway, or community in sample i; *w_k_* represents the number of metabolites catalyzed by an enzyme (*w_k_* = 1 for metabolite normalized flow); *logCPM_k_* is the log-transformed counts per million of the gene that encodes for a given enzyme; *f_k_* is the flow of information determined by Infomap for the node, or reaction *k*. The denominator of the equation normalized the weighted sum allowing the prevention of bias due to variations in the number of metabolites associated with the nodes. We assigned a value of 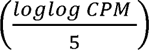 to reactions with no expression value in each sample to keep the metabolic network consistent across ovarian follicle development and reduce unconnected portions of the network. We assessed differences in normalized flow across different cell types and stages using the Mann-Whitney U test with False Discovery Rate (FDR) to correct for multiple comparisons and deemed them significant if their FDR < 0.05 ^35,36^. Feature ranking was based on the following order: FRD-corrected p-values (lower to higher) coefficient of variation (higher to lower), and number of comparisons that each feature was statistically significant (FRD-corrected p-values<0.05). To identify features consistently significant in a specific cell and stage model, we identified the number of times the feature was significant in a comparison (FDR < 0.0001) involving said cell and stage model and ranked them by the number of significance and then the logFC of the flow value to rank ties. To avoid false positives, we removed pathways with low biological confidence (Recon3D confidence score of 0 or 1) from these rankings. For communities, gene ontology enrichment was conducted using ClusterProfiler to identify biological processes and molecular functions associated with the enzymes in each community, beyond the predefined Recon3D subsystems^37^.

### Secreted and consumed metabolites identification

We calculated the normalized flow of information for each secreted and consumed metabolite in each sample (eq. 7) by considering the sign of the flux associated with each extracellular exchange reaction computed by FBA results for each model (−1, 0, 1).

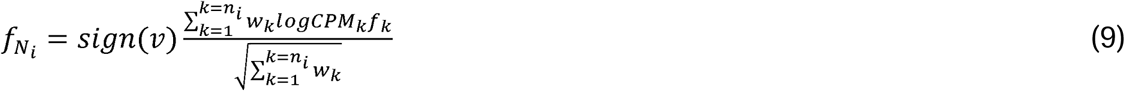

## RESULTS

### Computational pipeline to develop sample-specific context-informed metabolic networks

We developed a novel pipeline to generate sample-specific and context-informed genome-scale metabolic networks that account not only for the predicted enzymatic activity of their reactions, but also their local relationships (connectivity) (**Fig.1**). First, a weighted and directed network is generated from the stoichiometric matrix (S) of a context-specific GEM of interest, created using common approaches, like fastcore, IMAT, ftINT, in which the network nodes represent reactions, and the edges the metabolites shared between them. Following Beguerisse-Díaz and colleagues, we calculate the importance of each edge, *D_ij_*, between reaction *i* and reaction *j* as the summation of the probabilities that each metabolite present in the GEM, *k* = 1,2,3,..,*n*, can be produced by reaction *i* and consumed by reaction *j*

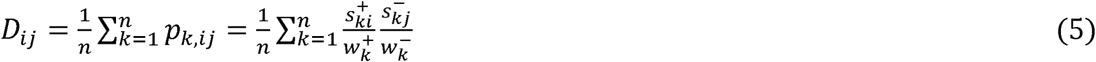

Therefore, we not only consider the contribution of each reaction for each metabolite with respect to all the reactions that it partakes in as a reactant or product, but also the reversibility of those reactions.

Departing from Beguerisse-Díaz and colleagues, instead of using the reaction flux to estimate reaction importance, we determine the importance of each node (reaction) within the metabolic weighted and directed network based on the flow of information between reactions (nodes) by minimizing the physics-derived map equation^34^. Briefly, the map equation determines the flow pattern of random walker based on its probability to remain in a region of the network (e.g., metabolic subsystem) versus moving to a different section of the network. Minimization of the map equation enables the identification of highly interconnected nodes, (modularity or communities) and the flow of information between nodes (reactions). Based on the flow of information within the context-specific weighted and directed metabolic network, we can determine the flow of information for a sample-specific weighted and directed metabolic network as the normalized flow of information for an enzyme-encoding gene, *i*, *f_Ni_*, based on the transcriptional activity of each enzyme-encoding gene *(logCPM_i_*) and their associated flow of information for each reaction it partakes in, *f_k_*, as an enzyme can catalyze multiple reactions, *n_i_*:

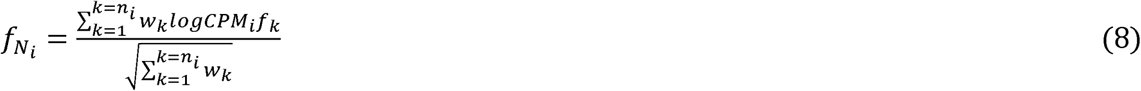

where *w_k_* represents the number of metabolites catalyzed by the enzyme in reaction *k*. The normalized flow of information accounts not only for the level of expression of each gene (proxy for their enzymatic activity) but also for its importance within the network based on its network connectivity. Thus, the normalized flow of information of enzyme-encoding genes between different samples can be statistically compared to identify differences during complex multi-cellular developmental processes at a systems level. Importantly, the concept can be extended to determine the normalized flow of information of metabolic pathways, internally consumed and secreted metabolites, and data-driven metabolic communities or modules (clusters of nodes).

### Sample-specific metabolic flow to understand human ovarian follicle development

To show the insights that our novel approach can provide, we used human ovarian follicle development as a model system. Ovarian follicle development is a dynamic and highly orchestrated multi-cellular processes in which metabolism is key, culminating in the production of a high-quality oocyte, the female germ cell, which is required for successful fertilization and early embryonic development **(Fig. 2)**^15^. The oocyte must accumulate the necessary metabolites to support meiotic divisions upon fertilization and sustain development until implantation in the uterine walls^38–40^.

First, we generated a human genome-scale ovarian follicle metabolic model by overlaying genes that encode for enzymes from publicly available human transcriptomics data (153 bulk RNA-seq samples, **Table S5**) with the latest human genome metabolic model, Recon3D. The human ovarian follicle GEM contained 5,494 metabolites, 3,031 genes, and 8,421 reactions, was able to pass 432 out of 481 metabolic functions and could be divided into 104 metabolic communities (or cluster of reactions)^16,34^. Using this human ovarian follicle GEM as the global model, we generated a total of 8 cell- and stage-specific metabolic models (2 cell types, the oocyte and granulosa cells; 4 stages of ovarian follicle development, primordial, primary, secondary and antral) by overlaying genes that encode from enzymes in each cell and follicle developmental stage **(Fig. 3)**. These context-specific models were used to determine the sample-specific metabolic flow of information for the total 153 samples available in the dataset (77 oocyte, 53 granulosa).

**Figure 3.**
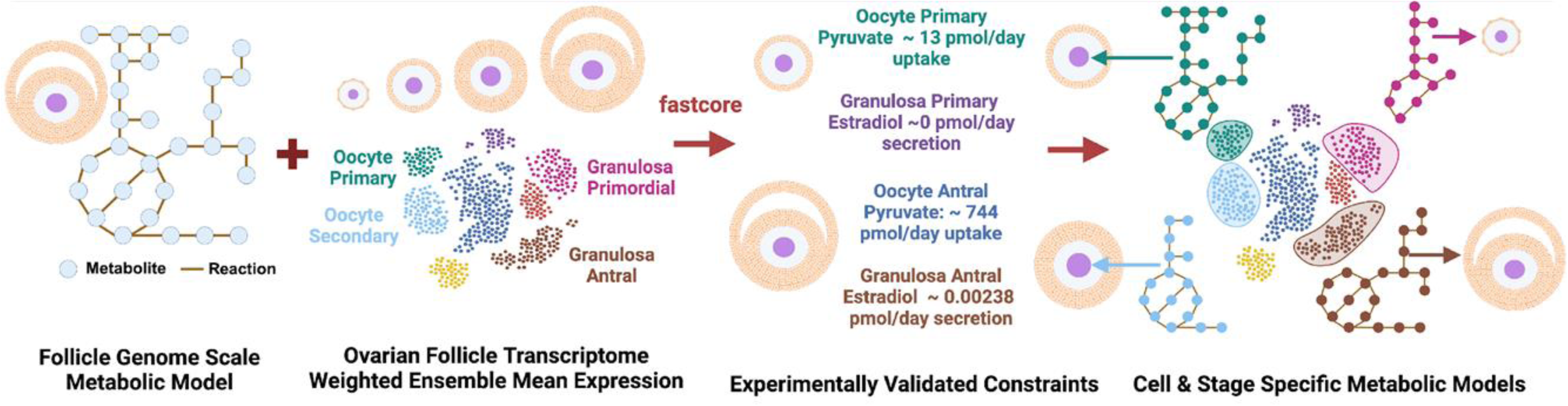
Methods for Generating GEMs of Ovarian Follicle Development. GEMs were developed by integrating ovarian follicle transcriptome data with Human Recon3D (fastcore). Enzymes were included in the model if their Weighted Ensemble Mean Expression > log(5). Cell and stage-specific models were constrained using experimentally validated values from literature.

Multiple of the 104 distinct communities identified by Infomap in the human ovarian follicle GEM, which varied in term of size and proportions of metabolic subsystems **(Fig. 4)**, changed their normalized flow of information throughout development. A total of 101 metabolic communities had a statistically significant different flow of information in at least one comparison between the different cells and developmental stages based on the sample-specific metabolic flow of information **(Table 1)**, with clear differences in the communities of the oocyte and granulosa cells across all stages of development and at different stages of development within each cell type. Oocytes and granulosa cells in primordial versus primary follicles presented one of the largest differences (75 and 83 communities, respectively) as well as between the oocytes and granulosa cells in primordial and primary follicles (83 and 88 communities, respectively). The primordial follicles were the least metabolically active (low normalized flow of information) compared with other stages, in agreement with a dormant stage; oocytes increased their metabolic activity between the primordial to primary transition while the granulosa cells delayed an increase in metabolic activity until the secondary to antral stage (**Fig. 5**). Among the top three communities that presented statistically distinct normalized flow across comparisons (communities 41, 58, and 11), their normalized flow in the oocyte was higher than in the granulosa cells for the same developmental stage, yet those differences were reduced as the ovarian follicle develops. (**Fig. 5, Table S6**). For instance, community 41, which included a total of 58 reactions, was composed of enzymatic processes belonging to ubiquinone synthesis (32.6%), glycine, serine, alanine, and threonine metabolism (15.2%) and cholesterol metabolism, fatty acid oxidation, and squalene and cholesterol synthesis (all 13.0%) and was enriched in energy and lipid metabolic processes, such as ubiquinone, cholesterol, steroid, and phospholipid biosynthesis (FDR < 0.005). Community 58 (43 reactions) was composed mostly of processes related with fatty acid oxidation and synthesis (56.1% and 29.3%, respectively) and fatty acid biosynthetic process and fatty acid beta-oxidation were enriched (FDR < 0.005), in agreement with prior results indicating that fatty acids are produced by the oocyte that highly expressed lipid beta oxidation enzymes ^41,42^. Finally, community 11 (144 reactions) was also composed by reactions associated with fatty acid oxidation (28.29%) and valine, leucine, and isoleucine metabolism (21.71%), with enriched process associated with fatty acid catabolism, the tricarboxylic acid cycle, and pyruvate metabolic process (FDR < 5e-08).

**Figure 4.**
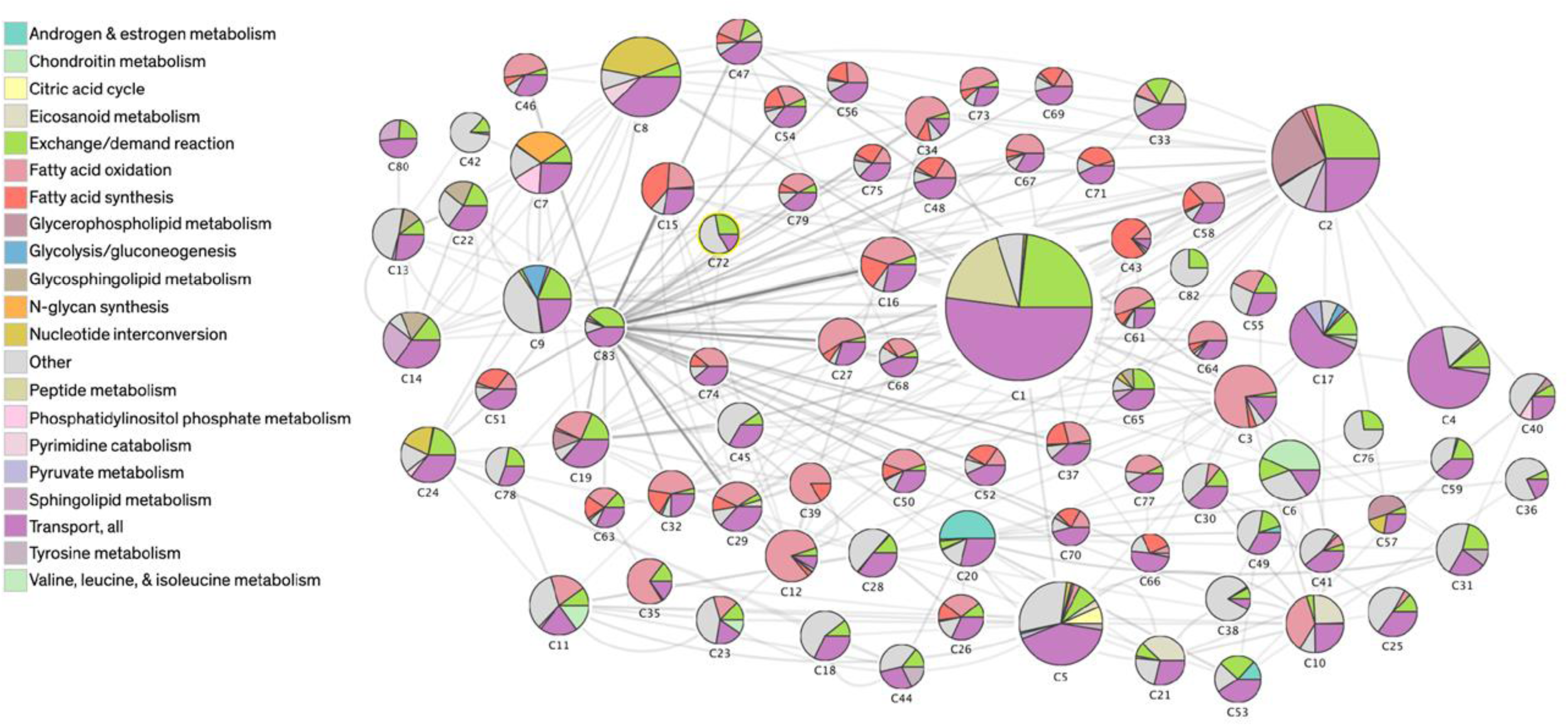
Several diverse metabolic communities were identified in the ovarian follicle genome-scale metabolic model and most of them included reactions belonging to different metabolic pathways. We employed Infomap for community detection in the ovarian follicle GEM. Colors represent the subsystems defined in Human Recon 3D (commonly canonical metabolic pathways) in which the reactions of each community are part of (we have limited to the top 20 pathways just for visualization purposes). The size of the divisions within each community corresponds to the proportion of reactions that are part of each subsystem. Size of the node corresponds to the number of reactions in the community. The thickness of the edge corresponds to the summation of flow of information from reactions that connect both communities.

**Figure 5.**
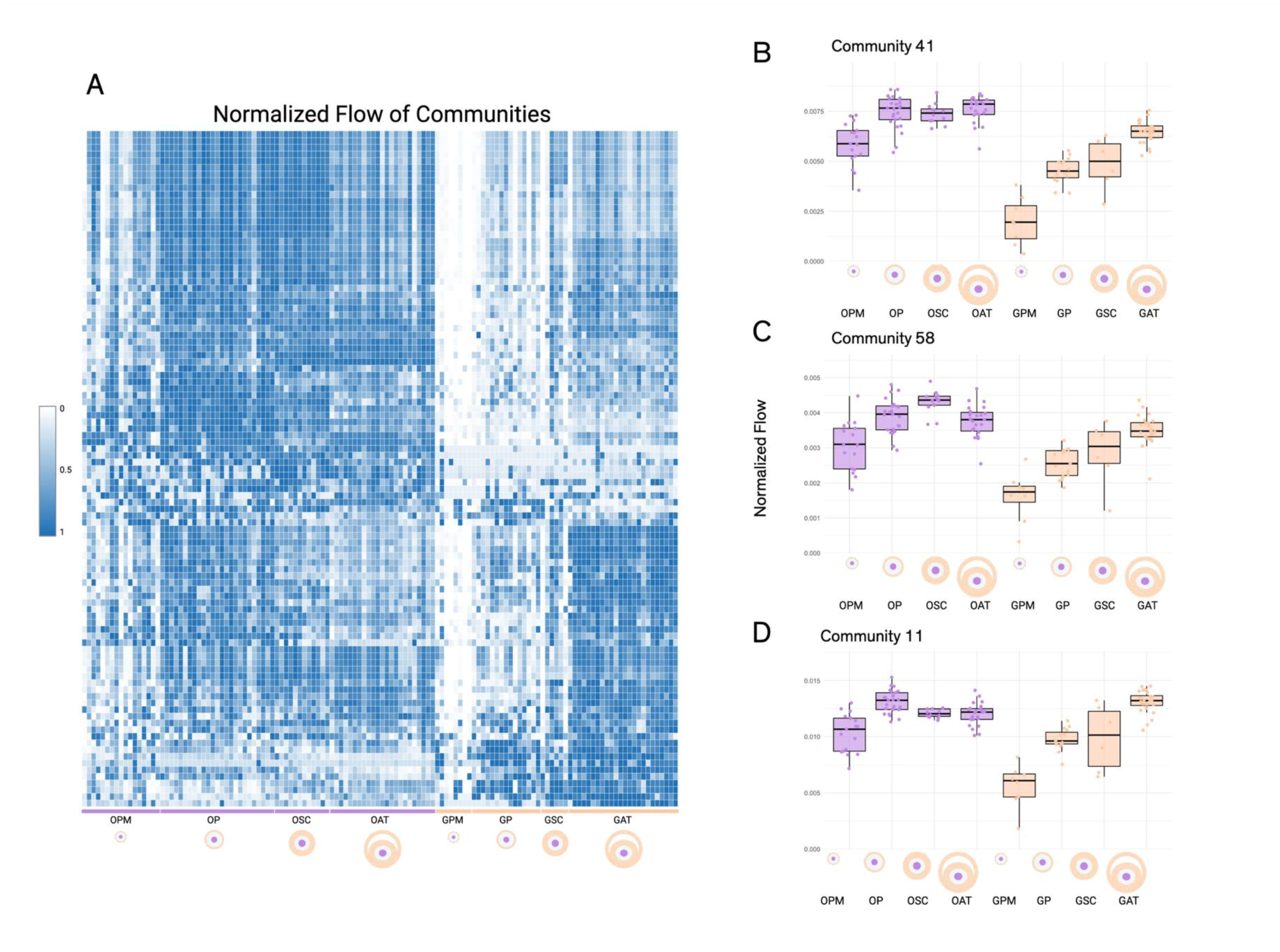
Sample Specific Normalized Flow through Follicle Communities. Communities were identified from the follicle GEM and sample-specific metabolic flow through each community was calculated by summing the normalized flow values of all nodes within each community per sample. A) Heat map of sample-specific normalized flow of all communities in each sample-specific metabolic models. Significant communities were determined by FDR values < 0.05 from Mann Whitney U Test and variance ratio to rank tie FDR values. The top three communities were plotted based on FDR and number of significant comparisons. B) Community 41 sample specific flow across all models. C) Community 58 sample specifc flow across all models. D) Community 11 sample specific flow across all models.

Our approach revealed the importance of several less studied canonical metabolic pathways or subsystems during ovarian follicle development. A total of 99 metabolic subsystems (out of 105) were statistically significant different in at least one comparison, most of them between antral granulosa cells vs antral oocytes (FDRs < 0.05, **Table 1**, **Fig. 6A**). Known metabolic pathways were found to be statistically significant such as pyruvate metabolism (granulosa versus oocytes in primary follicles, FDR = 2E-07) and androgen and estrogen metabolism (granulosa versus oocytes at the antral stage, FDR = 9E-09), as well as lesser studied in the context of ovarian follicular development. For instance, vitamin B6 metabolism within the oocyte was more significant in later stages of maturation (FDR = 5E-08), in contrast with the granulosa cells, in which the vitamin B6 metabolism had a higher normalized flow in primordial granulosa cells (FDR = 2E-04, **Fig. 6B**). Thiamine metabolism (vitamin B1) was significantly higher in the oocytes of early follicles compared with any other stages or the granulosa cells (FDR <1E-06, **Fig. 6C**). Another example is the glyoxylate and dicarboxylate metabolism that had high normalized flow in the oocyte compared to granulosa cells independently of the follicular stage, with a significant decrease in importance in granulosa cells as ovarian follicle mature (FDRs < 2E-03, **Fig. 6D**). When looking at metabolic pathways that were consistently significant for a specific cell and stage specific model, we identified the nucleotide salvage pathway (antral granulosa cells), heme degradation and chondroitin synthesis (primary oocytes), taurine and hypotaurine metabolism and starch and sucrose metabolism (antral oocytes).

**Figure 6.**
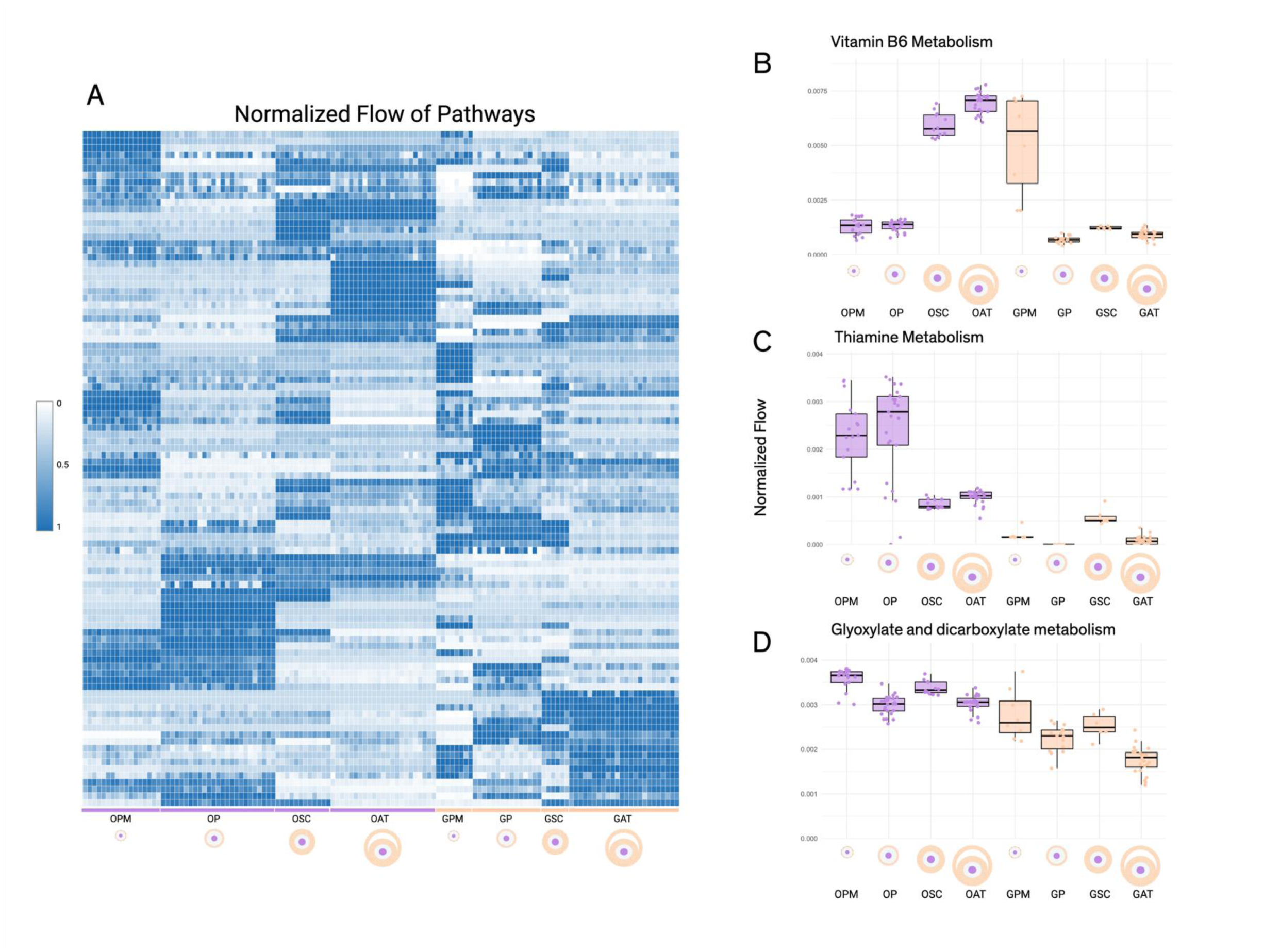
Sample Specific Normalized Flow through Metabolic Pathways. Pathways were determined based on the subsystem assigned to each reaction (node) by Recon3D. The flow through each pathway is calculated by summing the normalized flow values of all nodes within each pathway per sample. A) Heat map of sample-specific normalized flow of top significant pathways (top 25 per comparison) in each sample-specific metabolic models. Significant pathways were determined by FDR values < 0.05 from Mann Whitney U Test and variance ratio to rank tie FDR values. The top three enzymes were plotted based on FDR and number of significant comparisons. B) Vitamin B6 metabolism sample specific flow across all models. C) Thiamine Metabolism sample specific flow across all models; D) Glyoxylate and dicarboxylate metabolism sample specific flow across all

When exploring difference in normalized flow of information between enzyme-encoding genes, 2,618 unique genes had statistically significant different normalized flow across cell types and stages (FDR < 0.05, **Table 1**, **Figure 7A**), with most differences between the granulosa cells and the oocytes in antral follicles. The top three genes that encode for enzymes that were statistically significant in multiple comparisons were COX6A1 (cytochrome c oxidase subunit 6A1, FDRs < 0.005, **Fig. 7B**), HSD3B2 (hydroxy-delta-5-steroid dehydrogenase, 3 beta- steroid delta-isomerase 2, 5 comparisons with FDRs < 3E-05, **Fig. 7C**), and PLIN3 (perilipin 3, 3 comparisons with FDRs < 0.005, **Fig. 7D**). COX6A1 presented significantly higher normalized flow only in antral oocytes and in primary and secondary granulosa cells compared with the rest of conditions, while HSD3B2, a well-known steroid enzyme, was highest in granulosa cells after the secondary stage, in contrast with PLIN3, that was highest in the early stages of development of the granulosa cells (FDRs < 3E-07). We also observed cell-specific and stage-specific enzymatic markers, such as ATP2B1 and GLA for primary granulosa cells, GMPR and GMPR2 for antral granulosa cells, and FECH for primary oocytes.

**Figure 7.**
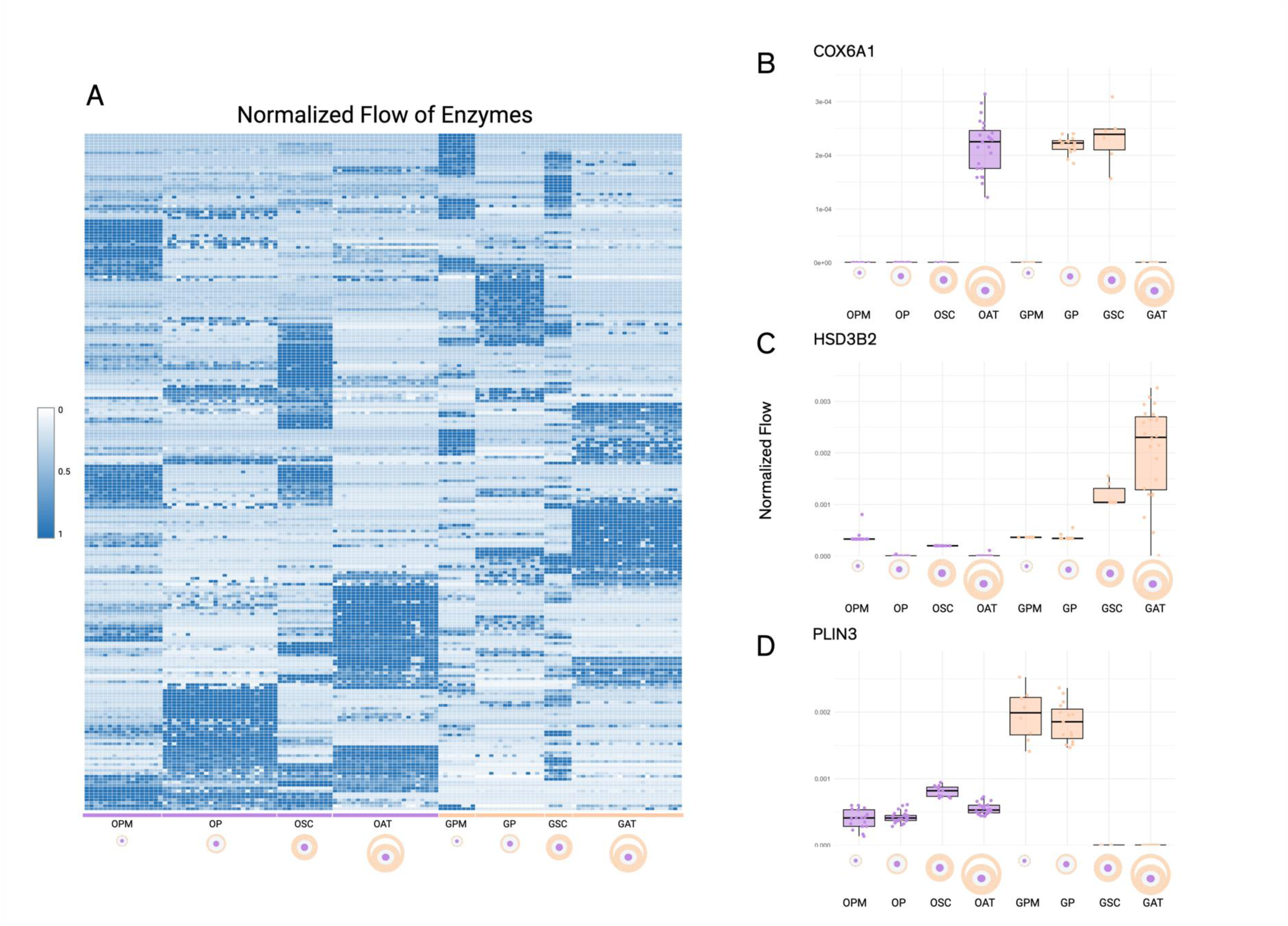
Sample Specific Normalized Flow of Enzymes. Enzymes were determined to be associated with a node based on the gene protein rules determined by Recon3D. A node’s flow was split equally among all enzymes associated with a node, and then the normalized flow for each enzyme was calculated. A) Heat map of sample-specific normalized flow of top significant enzymes (top 25 per comparison) in each sample-specific metabolic models. Significant enzymes were determined by FDR values < 0.05 from Mann Whitney U Test and variance ratio to rank tie FDR values. The top three enzymes were plotted based on FDR and number of significant comparisons. B) COX6A1 sample specific flow across all models. C) HSD3B2 sample specific flow across all models. D) RHBG sample specific flow across all mo dels.

Finally, we asked whether any metabolite changes their importance across cell type and stage. Similarly to the enzyme analysis, we identified several metabolites that had statistically different flow across cells and stages (**Fig. 8A**). Within all comparisons, there were a total 4,799 metabolites that were found to have a statistically significantly normalized flow across cells and stages with the most differences occurring between the antral stage between granulosa cells and the oocyte, and in the granulosa cells during antral formation (FDR < 0.05, **Table 1**). The top three most statistically significant metabolites across multiple models were all glycine derivatives that reside in the mitochondria, 3-methylcrotonoylglycine, isovalerylglycine, and 2-methylbutyrylglycine. They all follow the same trend with higher normalized flow in the oocytes during the primordial stage than in the secondary stage, granulosa cells presented opposite trends (FDRs < 1.5E-05, **Fig. 8B**). We also identified metabolites Our models also identified known metabolic differences including higher normalized flow of glucose in the primordial and antral stages in granulosa cells compared to the oocytes (FDRs<0.005). Interestingly, the most specific metabolites for a stage and cell type pertained to antral oocytes (FDR < 0.0001) as they were significant at many comparisons. These metabolites include the three mitochondrial metabolites 3α,7α - Dihydroxy-5β-cholestan-26-al, 3α,7 α,26-Trihydroxy-5β-cholestane, 2,4,7-decatrienoyl Coenzyme A, and in the nucleus, guanosine triphosphate and deoxycytidine triphosphate.

**Figure 8.**
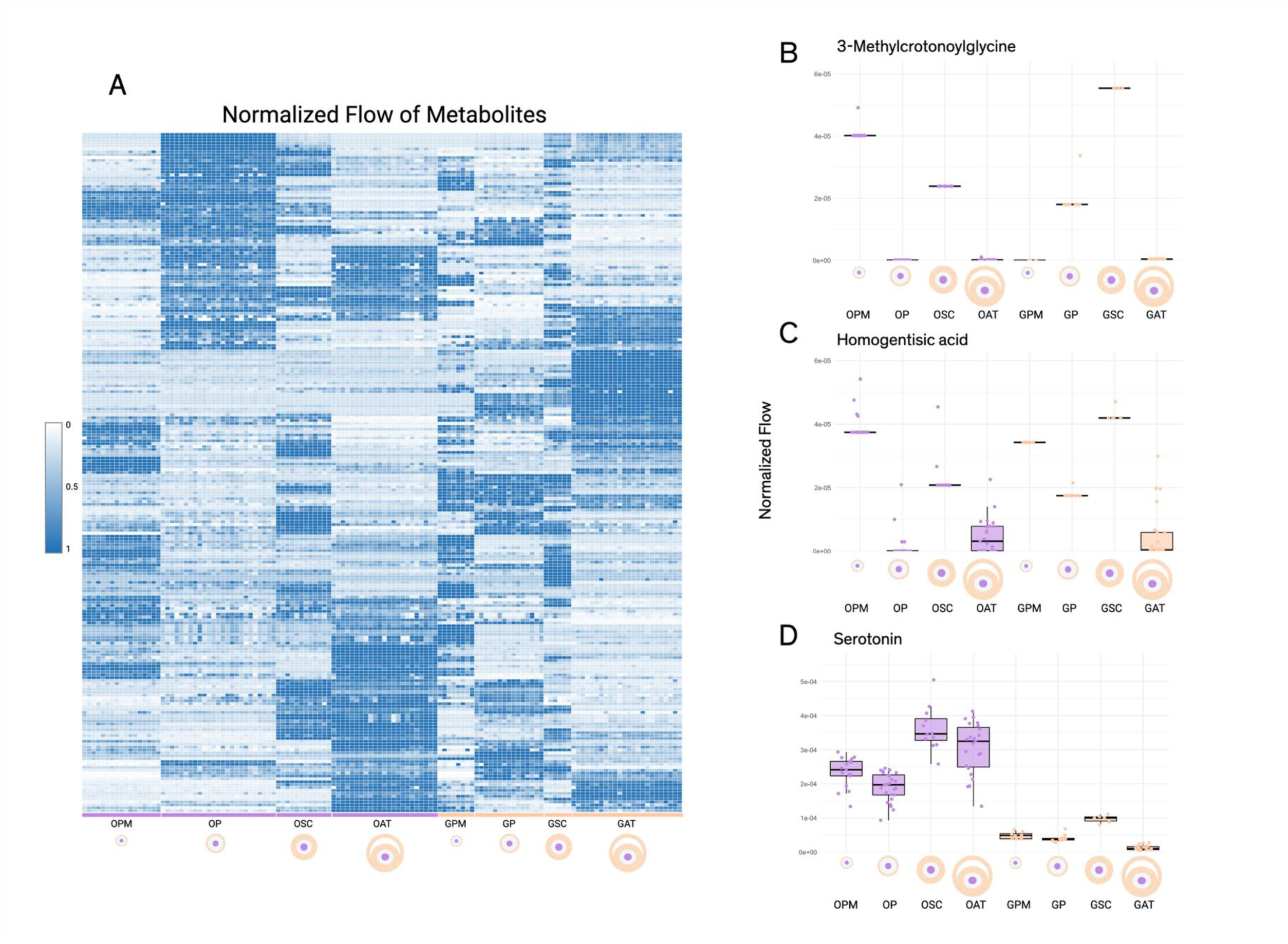
Sample Specific Normalized Flow of Metabolites. Metabolites were determined to be associated with a node based on the products and reactants of a reaction (node) as determined by Recon3D. A node’s flow was split equally among all netabolites associated with a node, and then the normalized flow for each metabolite was calculated. A) Heat map of sample-specific normalized flow of top significant metabolites (top 25 per comparison) in each sample-specific metabolic models. Significant metabolites were determined by FDR values < 0.05 from Mann Whitney U Test and variance ratio to rank tie FDR values. The top three metabolites were plotted based on FDR and number of significant comparisons. B) COX6A1 sample specific flow across all models. C) HSD3B2 sample specific flow across all models. D) RHBG sample specific flow across all models.

Possible translational applications of our new approach are the identification of predicted secreted and consumed metabolites during follicle development and the inter-cellular communication between the oocyte and granulosa cells **(Fig. 9)**. Out of the 250 statistically significant exchange metabolites (FDR < 5E-7 across any comparison), our results identify well-known metabolic activity, including the production of pyruvate by granulosa cells and consumption of it by the oocyte as well as the uptake of glucose by the granulosa cells^43,44^. Our models suggested an underexplored process during ovarian follicle development: the oocyte secretion of folic acid across all stages of development and by the granulosa cells at the primordial and secondary stages of development only. Interestingly, our results also predicted that folinic acid (vitamin B9), a derivative of folic acid, was consumed by the oocyte at all stages of development, but not the granulosa cells.

**Figure 9.**
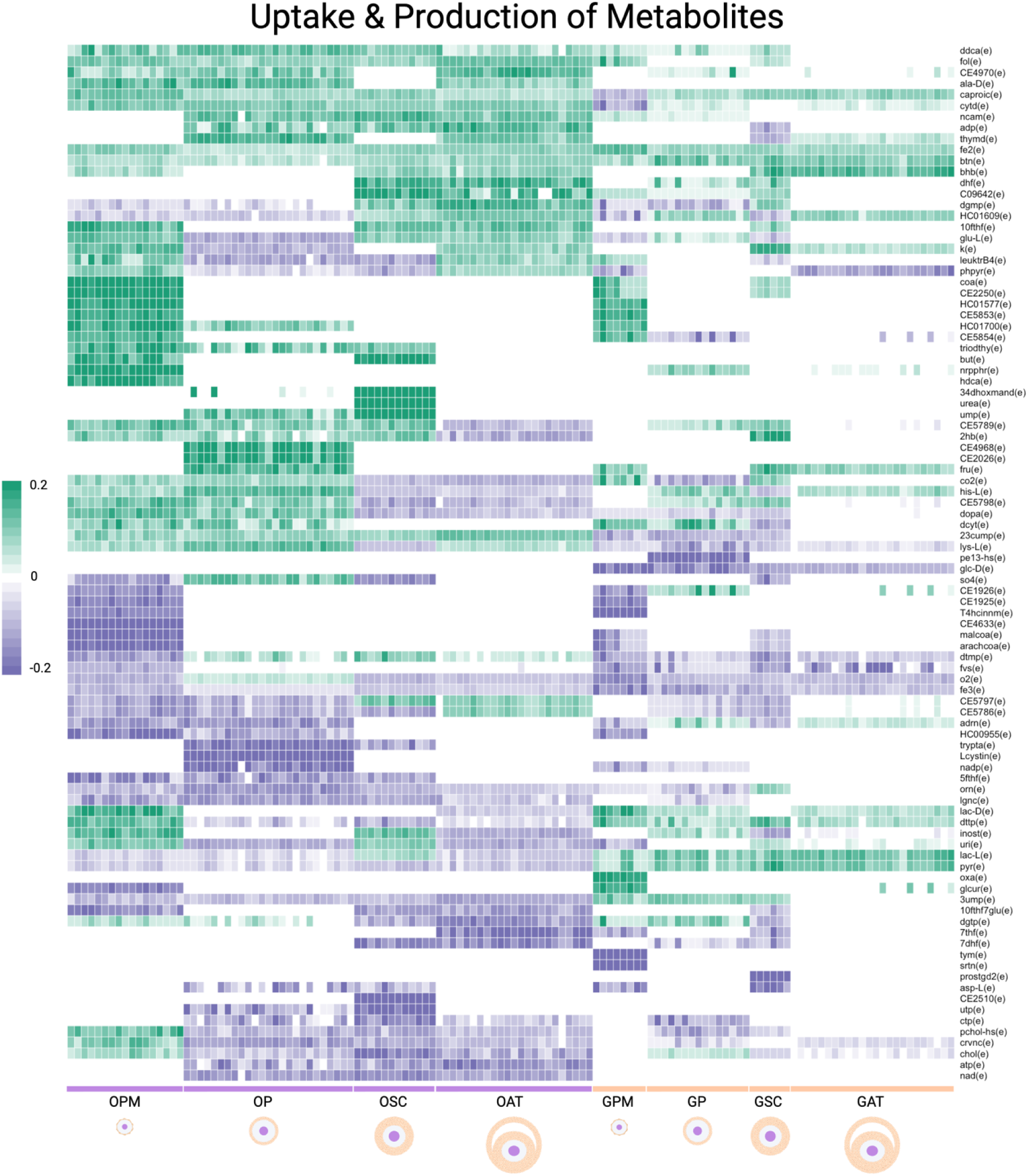
Sample Specific Normalized Flux. Top metabolites were determined based on sample-specific flow values. If this metabolite had an exchange reaction in the extracellular compartment, the normalized flow was multiplied by the flux balance anlaysis sign to indicate wether a metabolite is predicted to be consumed or produced by a given cell at a given stage.

## DISCUSSION

Here we presented a novel method that allows for the integration of multi-omics data and limited prior metabolic knowledge to render sample-specific and context-informed metabolic networks, enabling the study of dynamic metabolic processes in multi-cellular tissues. Our method naturally allows statistical comparisons across sample-specific and context-informed metabolic networks derived from GEMs, providing insights into how metabolic potential varies by cell types by accounting for sample (or individual) variability. Leveraging the statistical properties of the metabolic network structure allows the identification of the keystone metabolic communities, pathways, reactions, enzymes, and metabolites of inter-cellular and intra-cellular metabolic communications. Foget and colleagues proposed a similar approach using multi-organ genome-wide association studies (GWAS) data to create individual specific GEMs and subsequently calculate personalized organ-level fluxes with the goal of understanding the effect of genetic mutations into systemic metabolic levels but didn’t consider metabolic network connectivity^45^.

Importantly, while it is plausible to determine single-cell metabolic data, the technology is not yet ubiquitous, and our method offers a powerful framework for developing data-driven hypothesis to investigate the dynamics in understudied metabolic processes or conditions where limited information is known about their metabolism, yet transcriptomics data is available. Furthermore, and perhaps one of the main contributions of our work is that, compared with only transcriptomics and subsequent gene and pathway enrichment analysis, our approach integrates gene expression, manually curated metabolic reactions, and network structure, providing a systems-level view of metabolism, resulting in a deeper understanding of metabolic differences than just gene set enrichment analysis of metabolic processes based only on transcriptional levels.

To demonstrate the capability of our new approach, we focused on ovarian follicle development, a complex developmental process in which metabolism is key, yet the dynamic intricated inter-cellular and intra-cellular communications between the maturing germ-cell, the oocyte, and its neighboring supporting cells, granulosa cells, are mostly unknown. Our approach enables us, for the first time, to provide a broader perspective of the dynamic metabolism of ovarian follicles from metabolites, enzymes, subsystems and communities simultaneously in multiple cellular types. Based on the activities of the largest community within the metabolic networks, oocytes were more energetically demanding than granulosa cells across all maturation stages. This is consistent with the well-established role of energy and lipid metabolism in oocyte development^48^. Importantly, lipids and fatty acid metabolic processes emerged highly interconnected with energy production reactions, with coenzyme Qs (CoQ), acetyl-CoA acetyltransferase 2 (ACAT2), and solute carrier family 16 member 1 (SLC16A1) appearing within the same large metabolic communities, reflecting coordinated interactions between fatty acid oxidation, cholesterol metabolism and ubiquinone biosynthesis, respectively. Notably, the degree of cooperation among these processes was regulated by the maturity of the oocyte. While ACAT2 flow of information differed significantly across all developmental stages, SLC16A1 was only significantly different between primary and secondary oocytes, suggesting that the greatest demands for pyruvate and lactate import into the oocyte coincides with the primary to secondary transition, when granulosa cells are actively proliferating. In contrast, granulosa cells showed a delayed increase in overall metabolic activity, not rising until the secondary-to-antral transition, highlighting a striking asynchrony in metabolic activation between the two cell types. Together, these findings reveal stage-dependent coordination between fatty acid oxidation and cholesterol metabolism in the oocyte, and underscore the value of the community approach in uncovering metabolic dynamics that systems with limited data availability would miss.

Similarly, we can apply this systems-level perspective to predict dynamic metabolic processes at the pathway level. Among the top significant pathways was vitamin B6, which was significantly higher in granulosa primordial follicles and in oocytes of secondary and antral stages. This opposing pattern between cell types suggests a stage-dependent vitamin B6 utilization during folliculogenesis. In mouse models, vitamin B6 promotes primordial follicle activation through the PI3K/Akt signaling pathway in primoridal granulosa cells ^49^ and improves oocyte quality during in vitro maturation of oocyte-cumulus cells in bovine species^51^. Supporting this, pyridoxal kinase (PDXK), the rate limiting step enzyme for active vitamin B6 production^52^, was significantly higher in secondary versus primary oocytes (FDR = 7.14 E-6), suggesting that the demand for active vitamin B6 increases as the oocyte matures. From a more unbiased perspective, using our data-driven approaches, all reactions assigned to the vitamin B6 subsystem in the Human Recon3D model were grouped together in community 41 which also contains many reactions assigned to tryptophan, a precursor of serotonin, and tyrosine amino acid metabolisms, and tetrahydrobiopterin, an essential cofactor for the production of serotonin, dopamine and nitric oxide. The neurotransmitter serotonin can be generated locally by the ovarian follicle in the presence of vitamin B6 and tetrahydrobiopterin and further regulates progesterone production. Nitric oxide within the oocyte is essential for maintaining oocyte quality, and their production is reduced with aging due to lower levels of tetrahydrobiopterin. Taken together, while the biochemical connections between vitamin B6, tryptophan, and tetrahydrobiopterin through serotonin synthesis are established, their coordinated cell type and stage-specific activity within the ovarian follicle represents a previously unrecognized biological context that warrants direct experimental investigation.

COX6A1 presented its highest activity in the antral oocytes and in primary and secondary granulosa cells. Previous research indicating that COX6A1 is active in the murine oocytes to regulate the mitochondrial electron transport chain^53^. Similarly, our results capture the importance of HSD3B2, an essential enzyme to produce progesterone in the Ä4 pathway in the granulosa cells at later follicular stages. PLIN3, a lipid droplet-associated protein formation^54,55^ predominantly active in early folliculogenesis, in both oocytes and granulosa cells, is consistent with reports of declining PLIN3 expression as oocytes progress toward maturation. Importantly, we also revealed unappreciated roles of several enzymes during ovarian follicle development. For example, GLA was more significant in granulosa primordial metabolic models in the lipid regulation of early-stage follicles. While GLA has been studied as a modulator to reduce systemic symptoms of PCOS such as hyperandrogenism and inflammation, these studies have not been extended to the context of ovarian follicle development^56^.

At the metabolite level, multiple glycine-derived metabolites, such as 3-methylcrotonoylglycine, isovalerylglycine and 2-methylbutyrylglycine, were highly significant in the mitochondria of both oocyte and granulosa cells in early follicles. This is consistent with community 11 identified in the network analyses **(Fig 5),** which grouped fatty acid oxidation with valine, leucine, and isoleucine metabolism and enriched TCA cycle processes, together suggesting that active branched-chain amino acids (BCAA)-driven mitochondrial energy metabolism may be a feature of early folliculogenesis in both the oocyte and granulosa cells. N-isovalerylglycine has been identifies as a consistently significan metabolite across ovarian developmental stages in goats, while glycine has been reported to promote in oocyte maturation in pigs by regulating lipid peroxidation^58,59^. Together, this suggests a broader rolse for glycine metabolism in follicular competence across species. This analysis also identified homogentisic acid, a tyrosine catabolism intermediate, as significant in both cell types during the primordial and secondary stages. Its significance alongside the BCAA-derived acylglycines suggest that both, aromatic and BCAA catabolism are active in early follicles. This points to amino acid turnover as a metabolic feature of early folliculalogenesis.

Serotonin, was more significant in oocytes compared to granulosa cells, following the same trend and the tryptophan metabolism precursor in community 41. Further, several metabolites involved in bile acid biosynthesis were consistently significant (FDR < 0.0001) in comparisons involving the oocytes in antral models. The intermediary 3-alpha, 7-alpha-dihydroxy-5-beta-cholestan-26-al, and 3-alpha,7-alpha,26-trihydroxy-5beta-cholestane, are part of the primary bile acid biosynthesis pathway and were significant (FDR < 0.0001) in comparisons involving oocyte antral models. Bile acid metabolism is active in the human ovarian follicle and plays a key role in sterogenesis^60^. Bile acids are present in the follicular fluid of the human and bovine ovaries and oocytes and granulosa cells contain bile acid importers and exporters suggesting bile acids might have reached the follicular fluid by passive and active transport from blood^61^. Yet, based on our models it is plausible that 3 alpha,7 alpha,26-trihydroxy-5beta-cholestane might be secreted by the oocyte and the granulosa cells at all stages of development. While our results should be validated experimentally, our results demonstrate the value of integrating prior metabolic knowledge and omics data to provide a new level of understanding at the specific sample level.

Finally, our approach can also be employed to identify circulatory biomarkers to study dynamic processes, such as secreted metabolites, or to identify putative metabolites that can render specific outcomes, which could later be translated into clinical applications. For instance, we predicted that the oocyte and granulosa cells secrete folic acid and consume folinic acid (the metabolic active and reduced form of folic acid) since early folliculogenesis. Our prior results in murine ovarian follicle genome-scale metabolic models also suggested that the oocyte produced folic acid^6^. Yet, the role of folic acid during folliculogenesis has been limited to ovulation and fertilization rate. For instance, in humans, higher levels of folic acid were associated with adequate cyclicity and higher ovarian reserve^62,63^ and in animal models, addition of folic acid to the IVF media improves oocyte maturation and blastocyst development rate^64^. Furthermore, no published study up to date has explored the effects of folinic acid in the oocyte, yet folinic acid might directly or indirectly participate in critical oocyte process, such as DNA methylation of the growing oocytes, DNA repair mechanisms, buffering reactive oxidative species, or accumulation of folate for later divisions after fertilization. In summary, our approach uncovered significant metabolic features that drive follicular development from a systems level approach. These results provide a foundation for a broader exploration of mostly underexplored metabolic cooperation between the oocyte and granulosa cells throughout the distinct stages of development.

### Strengths and Limitations

Unlike transcriptomics alone, our approach integrates cell- and sample-specific gene expression, manually curated metabolic reactions, and network structure, providing a systems-level view of metabolism, resulting in a deeper understanding of metabolic differences than just gene set enrichment analysis of metabolic processes based only on transcriptional levels. Importantly, our systems-level approach generates data-driven novel hypotheses to understand the metabolism of complex tissues, and developmental processes, from community to metabolite levels, by taking advantage of commonplace omics data, such as bulk RNA-seq. In the context of women’s reproduction, most human studies on ovarian follicle development have focused on gene regulation, leaving metabolic activity largely unexplored^46,47^. Here, for the first time, we have leveraged RNA-seq data to generate cell and stage specific metabolic models of human ovarian follicles to study temporal metabolic dynamics.

There are multiple areas where our approach can be further improved. Insights gained using our approach are limited by the available information in the literature, yet the models can be updated as new information becomes available. Currently, transcriptional abundance is employed as a proxy for metabolic activity, but other biological measurements more closely related to enzymatic activity could be also utilized, such as proteomics, metabolomics, and lipidomics. Similarly, model reconstruction methods continue to be improved and can be further refined, by including thermodynamic and kinetic information. Moving from undirected metabolic networks towards bipartite weighted and directed metabolic networks that include as nodes both reactions and metabolomics could better represent the flow of metabolic information in the cell, and we plan to include them in our future work. Finally, while our approach is based on bulk transcriptomics data, it is possible to extend it to single-cell RNA-seq after accounting for the sparsity and other nuances of these datasets.

## Conclusion

The development of systems biology approaches that capture dynamic and cell-specific metabolic processes across developmental stages is critical for understanding the metabolic mechanisms in complex biological processes. In this study, we have applied our network-based method to the developmental context of ovarian folliculogenesis. Utilizing GEMs for two distinct cell types across four stages of development, we have generated data-driven hypotheses of metabolic activity, which can provide novel insights into this complex and dynamic process.

Metabolic understanding is not only important relevant for fundamental biology, but also for addressing multiple human conditions, such as common and poorly understood women specific disorders that occur during their reproductive lifespan, such as polycystic ovarian syndrome, premature ovarian insufficiency, and early menopause^65,66^. Overall, by developing context-aware, network-based approaches, we can generate extensive and testable hypotheses for other understudied biological systems, allowing for more targeted experimental studies.

## Data availability

GEO: GSE107746

Scripts will be made available on GitHub upon publication

## Contributions

BPB conceived the original idea and secured funding; EL developed all the models and computational approaches and performed all the subsequent analysis; EL and BPB interpreted the results and wrote the original manuscripts; and all the authors critically reviewed the manuscript. We would like to acknowledge, Dr. Yan, for their assistance with extracting the data used in this study.

## SUPPLEMENTAL INFORMATION

Table S1: Exchange reaction constraints based on plasma levels obtained from the Human Metabolome database.

Table S2: Predicted number of granulosa cells per stage of ovarian follicle development. Values were predicted based on literature.

Table S3: Metabolic reactions that were tested in each model to determine their functionality. A majority are previously published functions for Human Recon3D, with the additional ones being follicle specific reactions from previously published murine ovarian follicle models.

Table S4: Metabolic constraints for each cell- and stage-specific models based on literature.

Table S5: Number of cell type and stage samples per donor.

Table S6: Community composition, including the reactions (nodes) and the subystems they belong to according to Recon3D.

Table S7. Top 25 significant communities per cell and stage type comparison. For each comparison, the top 25 significant communities are recorded with their p-value, FDR, community ratio, and the ranking of communities based on the number of significant comparisons are included.

Table S8. All communities per cell and stage type comparison. For each community, the comparison, p-value, and FDR are recorded.

Table S9. Top 25 significant pathways per cell and stage type comparison. For each comparison, the top 25 significant pathways are recorded with their p-value, FDR, pathway ratio, and the ranking of pathways based on the number of significant comparisons are included.

Table S10. All pathways per cell and stage type comparison. For each pathway, the comparison, p-value, and FDR are recorded.

Table S11. Top 25 significant enzymes per cell and stage type comparison. For each comparison, the top 25 significant enzymes are recorded with their p-value, FDR, enzyme ratio, and the ranking of enzymes based on the number of significant comparisons are included.

Table S12. All enzymes per cell and stage type comparison. For each enzyme, the comparison, p-value, and FDR are recorded.

Table S13. Top 25 significant metabolites per cell and stage type comparison. For each comparison, the top 25 significant metabolites are recorded with their p-value, FDR, metabolite ratio, and the ranking of metabolites based on the number of significant comparisons are included.

Table S14. All metabolites per cell and stage type comparison. For each metabolite, the comparison, p-value, and FDR are recorded.

Table S15: Communities with an overrepresentation of significant results involving the same cell/stage model. For each community, the repeated model, number of significant comparisons, the comparisons, the logFC, FDR, and median and mean absolute logFC are included.

Table S16: Pathways with an overrepresentation of significant results involving the same cell/stage model. For each pathway, the repeated model, number of significant comparisons, the comparisons, the logFC, FDR, and median and mean absolute logFC are included.

Table S17: Enzymes with an overrepresentation of significant results involving the same cell/stage model. For each enzyme, the repeated model, number of significant comparisons, the comparisons, the logFC, FDR, and median and mean absolute logFC are included.

Table S18: Metabolites with an overrepresentation of significant results involving the same cell/stage model. For each metabolite, the repeated model, number of significant comparisons, the comparisons, the logFC, FDR, and median and mean absolute logFC are included.

## REFERENCES

1. Miyazawa H, Aulehla A. Revisiting the role of metabolism during development. Development. 2018;145(19):dev131110. doi: 10.1242/dev.131110. doi:10.1242/dev.131110

2. Passi A, Tibocha-Bonilla JD, Kumar M, Tec-Campos D, Zengler K, Zuniga C. Genome-Scale Metabolic Modeling Enables In-Depth Understanding of Big Data. Metabolites. 2021;12(1):14. doi: 10.3390/metabo12010014. doi:10.3390/metabo12010014

3. Ryu JY, Kim HU, Lee SY. Reconstruction of genome-scale human metabolic models using omics data. Integr Biol (Camb). 2015;7(8):859–868. doi:10.1039/c5ib00002e

4. Folger O, Jerby L, Frezza C, Gottlieb E, Ruppin E, Shlomi T. Predicting selective drug targets in cancer through metabolic networks. Mol Syst Biol. 2011;7:501. doi:10.1038/msb.2011.35

5. Foguet C, Xu Y, Ritchie SC, et al. Genetically personalised organ-specific metabolic models in health and disease. Nat Commun. 2022;13(1):7356–7. doi:10.1038/s41467-022-35017-7

6. Penalver Bernabe B, Thiele I, Galdones E, et al. Dynamic genome-scale cell-specific metabolic models reveal novel inter-cellular and intra-cellular metabolic communications during ovarian follicle development. BMC Bioinformatics. 2019;20(1):307–2. doi:10.1186/s12859-019-2825-2

7. Orth JD, Thiele I, Palsson BO. What is flux balance analysis? Nat Biotechnol. 2010;28(3):245–248. doi:10.1038/nbt.1614

8. Heinonen M, Osmala M, Mannerstrom H, et al. Bayesian metabolic flux analysis reveals intracellular flux couplings. Bioinformatics. 2019;35(14):i548–i557. doi:10.1093/bioinformatics/btz315

9. Raman K, Chandra N. Flux balance analysis of biological systems: applications and challenges. Briefings in Bioinformatics. 2009;10(4):435. doi:10.1093/bib/bbp011

10. Zhou W, Nakhleh L. Convergent evolution of modularity in metabolic networks through different community structures.

11. Ravasz E, Somera AL, Mongru DA, Oltvai ZN, Barabasi AL. Hierarchical organization of modularity in metabolic networks. Science. 2002;297(5586):1551–1555. doi:10.1126/science.1073374

12. Wunderlich Z, Mirny LA. Using the topology of metabolic networks to predict viability of mutant strains. Biophys J. 2006;91(6):2304–2311. doi:10.1529/biophysj.105.080572

13. Jeong H, Tombor B, Albert R, Oltvai ZN, Barabasi AL. The large-scale organization of metabolic networks. Nature. 2000;407(6804):651–654. doi:10.1038/35036627

14. Beguerisse-Diaz M, Bosque G, Oyarzun D, Pico J, Barahona M. Flux-dependent graphs for metabolic networks. NPJ Syst Biol Appl. 2018;4:32–y. eCollection 2018. doi:10.1038/s41540-018-0067-y

15. Fontana J, Martinkova S, Petr J, Zalmanova T, Trnka J. Metabolic cooperation in the ovarian follicle. Physiol Res. 2020;69(1):33–48. doi:10.33549/physiolres.934233

16. Brunk E, Sahoo S, Zielinski DC, et al. Recon3D enables a three-dimensional view of gene variation in human metabolism. Nat Biotechnol. 2018;36(3):272–281. doi:10.1038/nbt.4072

17. Zhang Y, Yan Z, Qin Q, et al. Transcriptome Landscape of Human Folliculogenesis Reveals Oocyte and Granulosa Cell Interactions. Mol Cell. 2018;72(6):1021–1034.e4. doi:10.1016/j.molcel.2018.10.029

18. Vanacker J, Camboni A, Dath C, et al. Enzymatic isolation of human primordial and primary ovarian follicles with Liberase DH: protocol for application in a clinical setting. Fertil Steril. 2011;96(2):379–383.e3. doi:10.1016/j.fertnstert.2011.05.075

19. Wang T, Yan L, Yan J, et al. Basic fibroblast growth factor promotes the development of human ovarian early follicles during growth in vitro. Hum Reprod. 2014;29(3):568–576. doi:10.1093/humrep/det465

20. Gougeon A. Dynamics of follicular growth in the human: a model from preliminary results. Hum Reprod. 1986;1(2):81–87. doi:10.1093/oxfordjournals.humrep.a136365

21. Bolger AM, Lohse M, Usadel B. Trimmomatic: a flexible trimmer for Illumina sequence data. Bioinformatics. 2014;30(15):2114–2120. doi:10.1093/bioinformatics/btu170

22. Dobin A, Davis CA, Schlesinger F, et al. STAR: ultrafast universal RNA-seq aligner. Bioinformatics. 2013;29(1):15–21. doi:10.1093/bioinformatics/bts635

23. Schneider VA, Graves-Lindsay T, Howe K, et al. Evaluation of GRCh38 and de novo haploid genome assemblies demonstrates the enduring quality of the reference assembly. Genome Res. 2017;27(5):849–864. doi:10.1101/gr.213611.116

24. Webster TH, Couse M, Grande BM, et al. Identifying, understanding, and correcting technical artifacts on the sex chromosomes in next-generation sequencing data. Gigascience. 2019;8(7):giz074. doi: 10.1093/gigascience/giz074. doi:10.1093/gigascience/giz074

25. Olney KC, Brotman SM, Andrews JP, Valverde-Vesling VA, Wilson MA. Reference genome and transcriptome informed by the sex chromosome complement of the sample increase ability to detect sex differences in gene expression from RNA-Seq data. Biol Sex Differ. 2020;11(1):42–9. doi:10.1186/s13293-020-00312-9

26. Anders S, Pyl PT, Huber W. HTSeq--a Python framework to work with high-throughput sequencing data. Bioinformatics. 2015;31(2):166–169. doi:10.1093/bioinformatics/btu638

27. Lewis JE, Forshaw TE, Boothman DA, Furdui CM, Kemp ML. Personalized Genome-Scale Metabolic Models Identify Targets of Redox Metabolism in Radiation-Resistant Tumors. Cell Syst. 2021;12(1):68–81.e11. doi:10.1016/j.cels.2020.12.001

28. Vlassis N, Pacheco MP, Sauter T. Fast reconstruction of compact context-specific metabolic network models. PLoS Comput Biol. 2014;10(1):e1003424. doi:10.1371/journal.pcbi.1003424

29. Jin S, Guerrero-Juarez CF, Zhang L, et al. Inference and analysis of cell-cell communication using CellChat. Nat Commun. 2021;12(1):1088–9. doi:10.1038/s41467-021-21246-9

30. Thiele I, Vlassis N, Fleming RMT. fastGapFill: efficient gap filling in metabolic networks. Bioinformatics. 2014;30(17):2529–2531. doi:10.1093/bioinformatics/btu321

31. Heirendt L, Arreckx S, Pfau T, et al. Creation and analysis of biochemical constraint-based models using the COBRA Toolbox v.3.0. Nat Protoc. 2019;14(3):639–702. doi:10.1038/s41596-018-0098-2

32. Wishart DS, Guo A, Oler E, et al. HMDB 5.0: the Human Metabolome Database for 2022. Nucleic Acids Res. 2022;50(D1):D622–D631. doi:10.1093/nar/gkab1062

33. Wishart DS, Tzur D, Knox C, et al. HMDB: the Human Metabolome Database. Nucleic Acids Res. 2007;35(Database issue):521. doi:10.1093/nar/gkl923

34. Rosvall M, Axelsson D, Bergstrom CT. The map equation. Eur Phys J Spec Top. 2009;178(1):13. doi:10.1140/epjst/e2010-01179-1

35. Mann HB, Whitney DR. On a Test of Whether one of Two Random Variables is Stochastically Larger than the Other. Ann Math Statist. 1947;18(1):50. doi:10.1214/aoms/1177730491

36. Benjamini Y, Hochberg Y. Controlling the False Discovery Rate: a Practical and Powerful Approach to Multiple Testing. 1995

37. Yu G, Wang L, Han Y, He Q. clusterProfiler: an R package for comparing biological themes among gene clusters. OMICS. 2012;16(5):284–287. doi:10.1089/omi.2011.0118

38. Conti M, Franciosi F. Acquisition of oocyte competence to develop as an embryo: integrated nuclear and cytoplasmic events. Hum Reprod Update. 2018;24(3):245–266. doi:10.1093/humupd/dmx040

39. Zuccotti M, Merico V, Cecconi S, Redi CA, Garagna S. What does it take to make a developmentally competent mammalian egg? Hum Reprod Update. 2011;17(4):525–540. doi:10.1093/humupd/dmr009

40. Krisher RL. The effect of oocyte quality on development. J Anim Sci. 2004;82 E-Suppl:14. doi:10.2527/2004.8213_supplE14x

41. Montjean D, Entezami F, Lichtblau I, Belloc S, Gurgan T, Menezo Y. Carnitine content in the follicular fluid and expression of the enzymes involved in beta oxidation in oocytes and cumulus cells. J Assist Reprod Genet. 2012;29(11):1221–1225. doi:10.1007/s10815-012-9855-2

42. Bao S, Yin T, Liu S. Ovarian aging: energy metabolism of oocytes. J Ovarian Res. 2024;17(1):118–y. doi:10.1186/s13048-024-01427-y

43. Sugiura K, Pendola FL, Eppig JJ. Oocyte control of metabolic cooperativity between oocytes and companion granulosa cells: energy metabolism. Dev Biol. 2005;279(1):20–30. doi:10.1016/j.ydbio.2004.11.027

44. Chiaratti MR, Garcia BM, Carvalho KF, et al. Oocyte mitochondria: role on fertility and disease transmission. Anim Reprod. 2018;15(3):231–238. doi:10.21451/1984-3143-AR2018-0069

45. Foguet C, Xu Y, Ritchie SC, et al. Genetically personalised organ-specific metabolic models in health and disease. Nat Commun. 2022;13(1):7356–7. doi:10.1038/s41467-022-35017-7

46. Bonnet A, Cabau C, Bouchez O, et al. An overview of gene expression dynamics during early ovarian folliculogenesis: specificity of follicular compartments and bi-directional dialog. BMC Genomics. 2013;14:904–904. doi:10.1186/1471-2164-14-904

47. Bonnet A, Servin B, Mulsant P, Mandon-Pepin B. Spatio-Temporal Gene Expression Profiling during In Vivo Early Ovarian Folliculogenesis: Integrated Transcriptomic Study and Molecular Signature of Early Follicular Growth. PLoS One. 2015;10(11):e0141482. doi:10.1371/journal.pone.0141482

48. Shi M, Sirard M. Metabolism of fatty acids in follicular cells, oocytes, and blastocysts. Reprod Fertil. 2022;3(2):R96–R108. doi:10.1530/RAF-21-0123

49. Wen J, Li W, Wang Z, et al. Vitamin B6 promotes the activation of primordial follicles through the PI3K/Akt signaling pathway. J Ovarian Res. 2025;18(1):296–1. doi:10.1186/s13048-025-01882-1

50. Zhang L, Wu L, Xu W, et al. Status of maternal serum B vitamins and pregnancy outcomes: New insights from in vitro fertilization and embryo transfer (IVF-ET) treatment. Front Nutr. 2022;9:962212. doi:10.3389/fnut.2022.962212

51. Aboelenain M, Balboula AZ, Kawahara M, et al. Pyridoxine supplementation during oocyte maturation improves the development and quality of bovine preimplantation embryos. Theriogenology. 2017;91:127–133. doi:10.1016/j.theriogenology.2016.12.022

52. Franco CN, Seabrook LJ, Nguyen ST, et al. Vitamin B(6) is governed by the local compartmentalization of metabolic enzymes during growth. Sci Adv. 2023;9(36):eadi2232. doi:10.1126/sciadv.adi2232

53. Feng Y, Wang J, Li M, et al. Impaired primordial follicle assembly in offspring ovaries from zearalenone-exposed mothers involves reduced mitochondrial activity and altered epigenetics in oocytes. Cell Mol Life Sci. 2022;79(5):258–0. doi:10.1007/s00018-022-04288-0

54. Zheng M, Andersen CY, Rasmussen FR, Cadenas J, Christensen ST, Mamsen LS. Expression of genes and enzymes involved in ovarian steroidogenesis in relation to human follicular development. Front Endocrinol (Lausanne). 2023;14:1268248. doi:10.3389/fendo.2023.1268248

55. Liu T, Qu J, Tian M, et al. Lipid Metabolic Process Involved in Oocyte Maturation During Folliculogenesis. Front Cell Dev Biol. 2022;10:806890. doi:10.3389/fcell.2022.806890

56. D Prabhu Y, Bhati M, Vellingiri B, Valsala Gopalakrishnan A. The effect of gamma-linolenic acid on Polycystic Ovary Syndrome associated Focal Segmental Glomerulosclerosis via TGF-beta pathway. Life Sci. 2021;276:119456. doi:10.1016/j.lfs.2021.119456

57. Zhang X, Zhang L, Xiang W. The impact of mitochondrial dysfunction on ovarian aging. J Transl Med. 2025;23(1):211–w. doi:10.1186/s12967-025-06223-w

58. Wang Y, Chao T, Li Q, He P, Zhang L, Wang J. Metabolomic and Transcriptomic Analyses Reveal the Potential Mechanisms of Dynamic Ovarian Development in Goats during Sexual Maturation. Int J Mol Sci. 2024;25(18):9898. doi: 10.3390/ijms25189898. doi:10.3390/ijms25189898

59. Gao L, Zhang C, Zheng Y, et al. Glycine regulates lipid peroxidation promoting porcine oocyte maturation and early embryonic development. J Anim Sci. 2023;101:10.1093/jas/skac425. doi:10.1093/jas/skac425

60. Smith LP, Nierstenhoefer M, Yoo SW, Penzias AS, Tobiasch E, Usheva A. The bile acid synthesis pathway is present and functional in the human ovary. PLoS One. 2009;4(10):e7333. doi:10.1371/journal.pone.0007333

61. Nagy RA, Hollema H, Andrei D, et al. The Origin of Follicular Bile Acids in the Human Ovary. Am J Pathol. 2019;189(10):2036–2045. doi:10.1016/j.ajpath.2019.06.011

62. Michels KA, Wactawski-Wende J, Mills JL, et al. Folate, homocysteine and the ovarian cycle among healthy regularly menstruating women. Hum Reprod. 2017;32(8):1743–1750. doi:10.1093/humrep/dex233

63. Kadir M, Hood RB, Minguez-Alarcon L, et al. Folate intake and ovarian reserve among women attending a fertility center. Fertil Steril. 2022;117(1):171–180. doi:10.1016/j.fertnstert.2021.09.037

64. Saini S, Sharma V, Ansari S, et al. Folate supplementation during oocyte maturation positively impacts the folate-methionine metabolism in pre-implantation embryos. Theriogenology. 2022;182:63–70. doi:10.1016/j.theriogenology.2022.01.024

65. Shelling AN, Ahmed Nasef N. The Role of Lifestyle and Dietary Factors in the Development of Premature Ovarian Insufficiency. Antioxidants (Basel). 2023;12(8):1601. doi: 10.3390/antiox12081601. doi:10.3390/antiox12081601

66. Chang RJ, Cook-Andersen H. Disordered follicle development. Mol Cell Endocrinol. 2013;373(1-2):51–60. doi:10.1016/j.mce.2012.07.011

